# Sex- and age-dependent effects of locus coeruleus-originated tau dysfunction on olfaction and neurophysiology in rats

**DOI:** 10.1101/2023.11.05.565692

**Authors:** Olivia D. E. Dutton, Marie A. Wasef, Alexander T. Burke, Ella A. Chirinos, Amelia T. Jones, Rachel Vey, Darlene M. Skinner, Carolyn W. Harley, Susan G. Walling

## Abstract

Braak and colleagues (2011) described human pretangle stages of abnormal tau protein originating in the locus coeruleus (LC) of young adults (Braak’s Stages a-c, 1a-b), decades prior to the neurofibrillary tangle stages observed in Alzheimer’s disease. To capture the features of LC-originated pretangle tau stages, we used a rat model where male and female TH-Cre^+/-^ rats received bilateral LC infusions of a hyperphosphorylated human tau (htauE14) gene via a viral vector. To assess the effects of age and sex on pretangle stage tau, we assayed physiological and behavioural changes 1-3mo (young) and 12^+^mo (aged) post LC-infusion (p.i.).

Open field measures revealed age and sex-dependent differences in anxiety in LC-htauE14 infused rats and overall sex- and age-dependent differences in activity. Odour discrimination tests showed little impairment in LC-htauE14 infused rats at 1-3mo after LC infusion. At 12^+^mo p.i. however, LC-htauE14 male rats failed a difficult odour discrimination test and a further olfactory detection test (habituation-dishabituation), but male LC-Control rats and female rats were not impaired.

LC neuronal firing rates were assessed in urethane-anesthetized aged (12^+^mo p.i.) male LC-htauE14 and LC-Control rats. The baseline level of firing of LC neurons in LC-htauE14 rats was higher than firing rates of LC-Control rats. Oscillatory patterns in LC firing were increased in amplitude and frequency by LC-htauE14.

PSD95 density measures in piriform cortex were similar for LC-Control and LC-htauE14 male and female rats of young and aged groups. While synaptophysin density was higher in controls of the 1-3mo p.i. groups than in the LC-htauE14 rats, levels did not differ among aged groups.

The increased excitability of LC cells observed may occur in relation to reduced LC axon arbours or may reflect a separate effect of htauE14 in LC neurons. The association of increased tonic LC activity to behavioural changes in the pretangle tau rat model merits further investigation.

## INTRODUCTION

The brainstem noradrenergic nucleus, the locus coeruleus (LC), has been identified as a potential site of origin of early-stage tau protein pathology (Braak et al., 2011; see Harley et al., 2021 for review). Traced first to the neurites of LC neurons of typically young adult human subjects in its earliest form (Braak’s Stage *a*), the soluble tau progresses through a hierarchical trajectory, first to the LC cellular compartment (Stage *b*) before transmission to other modulatory centers (e.g., raphe, and cholinergic regions) in Stage *c* prior to identification in regions in transentorhinal cortex (Braak’s Stage 1a and 1b). In contrast, Braak’s neurofibrillary tangle stages (Stage I-VI), are typically observed decades beyond the onset of pretangle tau dysfunction, making this pretangle stage an opportunistic period to identify early neuropathological and cognitive changes of Alzheimer’s disease (AD) in humans and animal models.

Prior to the manifestation of Alzheimer-related dementia and diagnosis, numerous neuropsychiatric and behavioural sensorineural changes have been identified as potential early markers of individual susceptibility for developing mild cognitive impairment (MCI) and AD. Of these potential early identifiers, notably the early loss of olfactory sensitivity, and/or the loss of the ability to identify olfactory stimuli is correlated with later cognitive decline and AD diagnosis (Devanand et al., 2015; Santabarbara et al., 2019). Pacyna et al. (2022) reported that a rapid decline in odour identification in aged individuals was highly associated with future MCI or AD diagnoses, even when they controlled for the presence of the APOE ε4 allele. Rapid olfactory decline was also associated with lower gray matter volume in olfactory and temporal regions (3T MRI). With reference to potential predictive neuropsychiatric symptoms, longitudinal analyses of the relationship between anxiety and future AD diagnosis in cognitively normal aged (Jang et al., 2020), or MCI progression to dementia, have indicated a relationship with anxiety scores and future AD risk (Jang et al., 2020; Cai et al., 2021). Despite evidence of the connection of the olfactory sensory deficits and anxiety-associated symptoms with future AD diagnoses, understandably few studies have examined these LC-related deficits in reference to Braak’s early pretangle stages of dysfunction in humans.

There have been two studies of pretangle LC-tau in TH-Cre rats using an adeno-associated virus (AAV) to introduce a modified human tau (htauE14) transgene that have successfully modelled Braak’s pretangle stage pathology (Ghosh et al., 2019; Omoluabi et al., 2021). Both studies have chronicled the progression of htauE14 from LC origins in male and female rats (sexes combined), documented noradrenergic fibre loss in forebrain regions (e.g., piriform cortex, PCx), and identified loss of olfactory discrimination and detection. Further, Omoluabi et al. identified increased anxiety in LC-htauE14 infused rats in one measure of anxiety (marble burying test), but not all anxiety-related tests (i.e., open field, elevated plus maze), 5mo post-LC-htauE14 infusion (p.i.).

The present study contrasts the behavioural and physiological effects of LC-directed AAV-htauE14 infusion performed in young adult TH-Cre^+/-^ male and female rats (3mo of age) at periods of 1-3mo (young) and 12^+^mo p.i. (aged). Tests of activity/anxiety and simple and difficult odour discrimination tasks are reported. Due to poor performance in odour discrimination and sensitivity tests in aged male LC-htauE14 rats, we recorded LC single unit activity in behaviourally naïve LC-htauE14 and LC-Control aged males. We found increased basal firing levels and increases in neural firing oscillation frequency in the LC-htauE14 infused rats compared to LC-Control rats. Lastly, we examined the pre- and post-synaptic markers synaptophysin and PSD-95, respectively, in the primary olfactory region, the PCx, and found early but transient changes in synaptic input, suggesting the documented loss of noradrenergic projections to this region reported previously (Ghosh et al., 2019; Omoloubi et al., 2021) is likely responsible for the changes in olfactory discrimination and detection in aged LC-htauE14 male rats.

## MATERIALS AND METHODS

### Subjects/breeding

Subjects were male and female TH-Cre^+/-^ rats bred at Memorial University using male TH-Cre^+/+^ (Sage Horizon) x female Sprague Dawley (Charles River, QC Canada) rats. After weaning, rats were housed with a regular 12:12h light-dark cycle (0700h-1900h), temperature (21-23°C), with two to three rats/cage in ventilated cages (Techniplast, CA), until 2-3mo of age. To reduce any confounding obesity-related health issues arising from age-associated obesity observed under conditions of *ad libitum* feeding, at the age of 2-3mo, both male and female rats were placed on a ‘moderate’ food restriction schedule, whereby rats received ^∼^75% of the ad libitum total food mass (Teklad2018), until the end of the procedures (see Hubert et al., 2000).

### LC-htauE14 Infusion Surgeries

The virus and transgenes infused were AAV2/9-Ef1α-DIO-eGFP-htauE14 (2.05e13 vg/mL; Virovek CA, USA; the htauE14 plasmid, a kind donation from Karen Ashe, Addgene), or AAV2/9-Ef1α-Flex-eGFP (1.8E13 vg/mL; Neurophotonics; Laval, QC CAN). Surgical procedures were modified slightly from those of Ghosh et al. (2019). In brief, at the age of 3mo, rats were anesthetized with isoflurane (2-5%; 1-2 L/min O_2_ flow rate), with meloxicam-SR analgesia (4mg/kg, 10mg/mL, s.c.) and placed in the skull flat position. Burr-holes were drilled bilaterally according to a sex- and size-dependent scale ranging from ^-^12.1mm to ^-^12.6mm posterior and ±1.3mm lateral from bregma. A 22ga stainless steel guide cannula (Plastics One, Anjou, QC) was lowered at a 20° angle to a final depth of ^-^6.4mm to ^-^6.7mm from brain surface to be situated in the LC-or immediate anterior dendrites. A separate infusion 28ga cannula extending 2.7mm from the bottom of the guide cannula was connected to a 5μL microsyringe (Hamilton) by autoanalyzer tubing (FisherBrand), and two 0.5μL infusions, separated by +100μm (lateral), over 2min were performed in each hemisphere (total 2μL volume/rat). After each infusion, the cannula was left in place for a minimum of 2min before slowly removing. After recovery (∼7 days), rats were transferred to a reversed light room (12:12h, lights off 0700h). Rats were allowed a minimum period of one month after surgery to allow LC-htauE14-eGFP expression before commencement of behavioural testing. Control conditions: sham infusion (cannula placement without infusion), control AAV-GFP infusion, or no surgery were combined to reduce animal numbers. Control AAV-GFP infused rats are identified in control condition data.

### Behavioural Testing

#### Activity and Anxiety -Open Field

Twenty-four hours prior to the start of the odour discrimination testing, rats were habituated to the testing apparatus (black plastic open field, 100cm x 100cm x 50cm height) for 10min. Anxiety and activity measures were analyzed using Noldus Ethovision14 (distance travelled, cm; total time in the center of maze, s; and total entries into the center) and Behavioral Observation Research Interactive Software (BORIS v7.13; number of supported and unsupported rears and total time spent grooming. For an animal to be considered in the center of the maze, the rat’s center point (Ethovision) had to cross the boundary of a square center area, measuring 60cm^2^ (adapted from Seibenhener & Wooten, 2015).

#### Odour Discrimination Task

##### General Odour Procedures

Two odour discrimination tasks were performed: a simple odour discrimination task (SOD; almond versus banana extract) and a difficult odour discrimination task (DOD; 0.001% 1-heptanol versus a 50:50 solution of 0.001% 1-heptanol and 0.001% 1-octanol in mineral oil). The experimental design and procedures were adapted from those of Tronel and Sara (2003). SOD and DOD tests were performed in the same open field apparatus which contained three sponges (8cm x 7cm x 2.5cm) with a hole (2cm in diameter) cut into the center. Sponges were placed in clear glass holders with the same approximate dimensions. In both tasks, each sponge was infused with 15μl of the odourants on each corner and placed in alternating corners of the open field box; one sponge was infused with the target odour (S^+^), and the other two sponges were infused with the non-target odour. The spatial configuration of the sponges alternated between trials to discourage rats from using associated spatial cues to locate the target sponge, and the target S^+^ odour was counterbalanced to be either one of the two paired scents but remained consistent for each animal during the tests. Both SOD and DOD tasks consisted of six trials, completed in one day. The DOD task began seven days after the completion of the SOD task.

#### Odour Discrimination Testing

On the first trial of the SOD and DOD tests, the target sponge was cued by placing Froot Loop^®^ pieces (approximately 1/3; the reward used throughout) on the top of the sponge. At the end of each trial, the sponges were removed and rinsed in odour-dedicated plastic containers containing ∼8L of tap water, using odour-dedicated gloves such that sponges were not contaminated with another scent. Odours were replenished at the beginning of each trial. Rats were trained to associate one of two odours with the reward in a scintillation vial cap placed at the bottom of the center hole of the sponges. During all tests, S^-^ (non-target) sponges were baited with an inaccessible reward (under the scintillation cap) to discourage rats from odour tracking. In the difficult task, rats were trained to differentiate between two similar odours (0.001% 1-heptanol versus a 50:50 solution of 0.001% 1-heptanol and 0.001%-octanol in mineral oil). Each trial lasted 5min or was terminated after the rat retrieved the reward. When the rat completed testing, the maze was wiped with 70% EtOH. After the first trial, the percentage of trials the rat chose to nose-poke the hole in the correct (S^+^) sponge, and latency to nose-poke the target sponge served as measures of performance. To facilitate learning of the discrimination task prior to commencement of the more similar odours in the DOD tests, the number of trials in the SOD test was extended to six in total. The first four trials were analyzed for each group. In analysis, a 120s cap was placed on latency measures.

#### Odour Detection Task (Habituation-Dishabituation)

To evaluate the ability to detect the odourants in aged rats, a four-trial habituation/dishabituation task was used. Each trial lasted 5min (n=5 each group). In each trial, two sponges were placed in opposite corners within the same open field apparatus used for odour testing. During Trials 1-3, both sponges were infused with 60μl of mineral oil (15μl on each corner). During the fourth trial, one sponge (the novel odour sponge) was replaced with sponge infused with 60μl of either 0.001% 1-heptanol made in mineral oil or the 50:50 solution of 0.001% 1-heptanol and 0.001% 1-octanol in mineral oil (DOD odours), or either the banana or almond extract (SOD odours). The second sponge was again infused with mineral oil, identical to the previous three trials. The position of the two sponges in the open field box was consistent across trials for each rat but was counterbalanced across rats. Percentage of time spent sniffing the novel sponge during Trial 4 (i.e., novel/dishabituation trial) was compared to that of Trial 3 (i.e., the last habituation trial). Rats were considered to be sniffing a sponge if they were within 2.5cm of the sponge and sniffing toward it. All four odours were tested and responses were averaged for presentation of SOD (banana and almond) and DOD (0.001% 1-heptanol and 0.001% 1-octanol) odour detection results.

### Locus coeruleus unit recording in LC-htauE14 infused and LC-Control aged male rats

A total of seven behaviourally naïve aged (12^+^mo post LC-infusion) male rats (four LC-htauE14, and three sham or non-operated control rats; ∼750g) were anesthetized with urethane (1.5g/kg, i.p) and placed in skull flat position in a stereotaxic instrument. Trephine holes were drilled bilaterally (3.8 – 4.2 AP; 1.2 - 1.4mm lateral to lambda; approximately 13.8-14.6 AP; 1.2 – 1.4mm lateral to bregma) to be situated over the LC and a 2-4 MΩ tungsten recording electrode (FHC Inc; UEWSGFSE4TNM) was lowered (∼5.9-6.6mm) at a 20° angle from cerebellar surface. LC units were collected using SciWorks 9.0 (Datawave) software, filtered 600Hz-10kHz, digitized 10kHz and stored for unit analysis.

#### LC unit analysis

During recording, LC units were identified with an audio analyser and oscilloscope based on characteristic low firing frequency, high amplitude with respect to baseline, and a phasic burst in response to toe-pinch. LC units were recorded with a tungsten microelectrode (4 MΩ; FHC, Bowdoin, ME, USA). The signal was amplified (Grass Instrument Co., Quincy, MA, USA) with a high-pass filter of 600Hz and low-pass of 10kHz. The signal was digitized with SciWorks 9.0 software (DataWave, Loveland, CO, USA) at 20kHz. Recorded units which reached an amplitude threshold of ≈1.5 × background were isolated for spike sorting. Spike sorting was performed manually with Datawave using a template matching strategy, emphasising spike amplitude, half-width, and slope (see Figure 5D) to sort waveforms into clusters. Clusters were then vetted based on amplitude, half-width, and slope averages, and those with low amplitude, gradual slope, or otherwise large variation from average unit parameters were excluded from further analysis. For each subject, 60s recordings were analysed for baseline LC firing frequency (Hz). A student’s t-test was performed to compare the firing frequency of LC-htauE14 and LC-Control subjects. A non-linear sign wave function was applied to the LC-htauE14 and LC-Control group total 60s baseline frequency data to examine macroscale changes in LC firing oscillations (Graphpad Prism; v9.5.1).

### Histology and Immunohistochemistry

Rats were anesthetized with urethane (1.5g/kg, i.p.), and either underwent in vivo electrophysiogical procedures (hippocampal field recordings, results not reported here), LC single unit recordings and then euthanized through transcardial perfusion (0.9% saline followed by buffered 4% paraformaldehyde; pH 7.4-7.6), or euthanized by urethane anesthetized perfusion without recording. After removal, brains were post-fixed in 4% paraformaldehyde overnight, then cryoprotected in 20% sucrose made in 0.1M PBS until sunk before being flash frozen and stored at ^-^80°C for further processing. Brains were sectioned at 30μm in both the coronal (brainstem) and horizontal (forebrain) planes. Forebrain sections were sampled throughout the dorsal-ventral axis with every ∼10^th^ section stained with cresyl violet (every ∼7 section for brainstem) and remaining sections were stored in long-term (polyvinylpyrillidone) storage solution in 24 or 48 well culture plates and placed in cold storage. In the case of LC recording, the brainstem was separated from forebrain prior to freezing, and subsequently sectioned in the coronal plane 20° from horizontal to verify electrode placement.

#### Immunohistochemistry

Free floating sections were processed for brightfield (diaminobenzidine; DAB) and fluorescence immunohistochemistry using a free-floating Tris buffered protocol, as previously described (Walling et al., 2007; Walling et al., 2012; Quinlan et al., 2019). Synaptophysin, and PSD-95 immunohistochemistry was performed on adjacent sections were processed using DAB-metal enhancement for brightfield microscopy. Tyrosine hydroxylase (TH), and green fluorescent protein were processed with immunohistofluorescence (Walling et al., 2012; Quinlan et al., 2019). To reduce the lipofuscin signal in aged animals i.e., 12mo^+^ p.i., TrueBlack™ lipofuscin quencher was used (0.5mL/section; 2min). See Table 1 for source and dilution information.

**Table 1.** Immunohistochemical sources and dilution information.

| Target | Host | Supplier | Product Number | Dilution |
| --- | --- | --- | --- | --- |
| **<u>Primary Antibodies</u>** |  |  |  |  |
| Tyrosine hydroxylase (TH) | Mouse | Millipore | MAB318 | 1:5,000-10,000 |
| Green Fluorescent Protein | Rabbit | ThermoFisher | A-11122 | 1:5,000 |
| PSD-95 | Mouse | ThermoFisher | MA1-045 | 1:2500 |
| Synaptophysin | Rabbit | ThermoFisher | PA527286 | 1:5000 |
| **<u>Secondary Antibodies</u>** |  |  |  |  |
| Alexa488 anti-rabbit | Goat | ThermoFisher | A-11034 | 1:400 |
| Alexa555 anti-mouse | Donkey | ThermoFisher | A-31570 | 1:400 |
| Biotinylated anti-mouse | Horse | Vector Labs | BA-2000 | 1:400 |
| Biotinylated anti-rabbit | Goat | Vector Labs | BA-1000 | 1:1,000 |
| **<u>Miscellaneous</u>** |  |  |  |  |
| DAB-metal kit |  | ThermoFisher | 34065 | As directed |
| TrueBlack (lipofuscin quencher) |  | Biotium | 23007 | As directed |

#### Image Analyses

Brightfield digital image acquisition was performed using an Olympus BX51 microscope and DP90 digital camera on LC (20x), and PCx (10x). Densitometry (relative optical density, ROD) was performed on PSD95 and synaptophysin immunolabelled sections in PCx using FIJI v2.1.0 on converted 8-bit grayscale images (5 circular samples, 75 pixels) on tissue sectioned in the horizontal plane. Fluorescent images were acquired with the Olympus BX51microscope with use of an epifluorescence light source (X-Cite Series 120).

LC neuron counts were performed on images of the Nissl-stained sections at 20x magnification of rats in the LC-htauE14 and LC-Control conditions in the LC recording experiment (aged 12^+^mo p.i.). Every 3^rd^ section through the LC was slide mounted and stained and a manual count was performed on identified neurons according to the criteria of Garcia-Cabezas et al. (2016). The average neuron count of the three sections containing the largest LC area was used and average number of neurons and average LC area per section are reported.

## RESULTS

To examine the behavioural effects of early age (3mo) LC-htauE14 infusion in male and female rats, we utilized two behavioural tests, designed to test activity and anxiety, and olfactory discrimination at two time periods post-LC infusion (young, 1-3mo p.i. and aged, 12^+^mo p.i.).

### Activity and Anxiety-Open Field Test (OFT)

Activity and anxiety variables (time in center, number of center entries, supported rears, unsupported rears, time spent grooming, and distance) were measured in male and female rats 1-3mo and 12^+^mo after LC-htauE14 or LC-Control infusions during the open field habituation session. The full 10min of the habituation test was analysed using a three-way (age, sex, LC-condition) factorial ANOVA. No three-way interactions were found for any variables (p>0.05), but interactions of sex x LC-condition, and age x LC-condition were revealed for center-focused variables. Across all six measured variables significant main effects of age, sex or sex x age interactions were also found (Figure 1). These results are presented relative to LC-condition, and then interactions or main effects relating to the age, and sex conditions.

**Figure 1.**
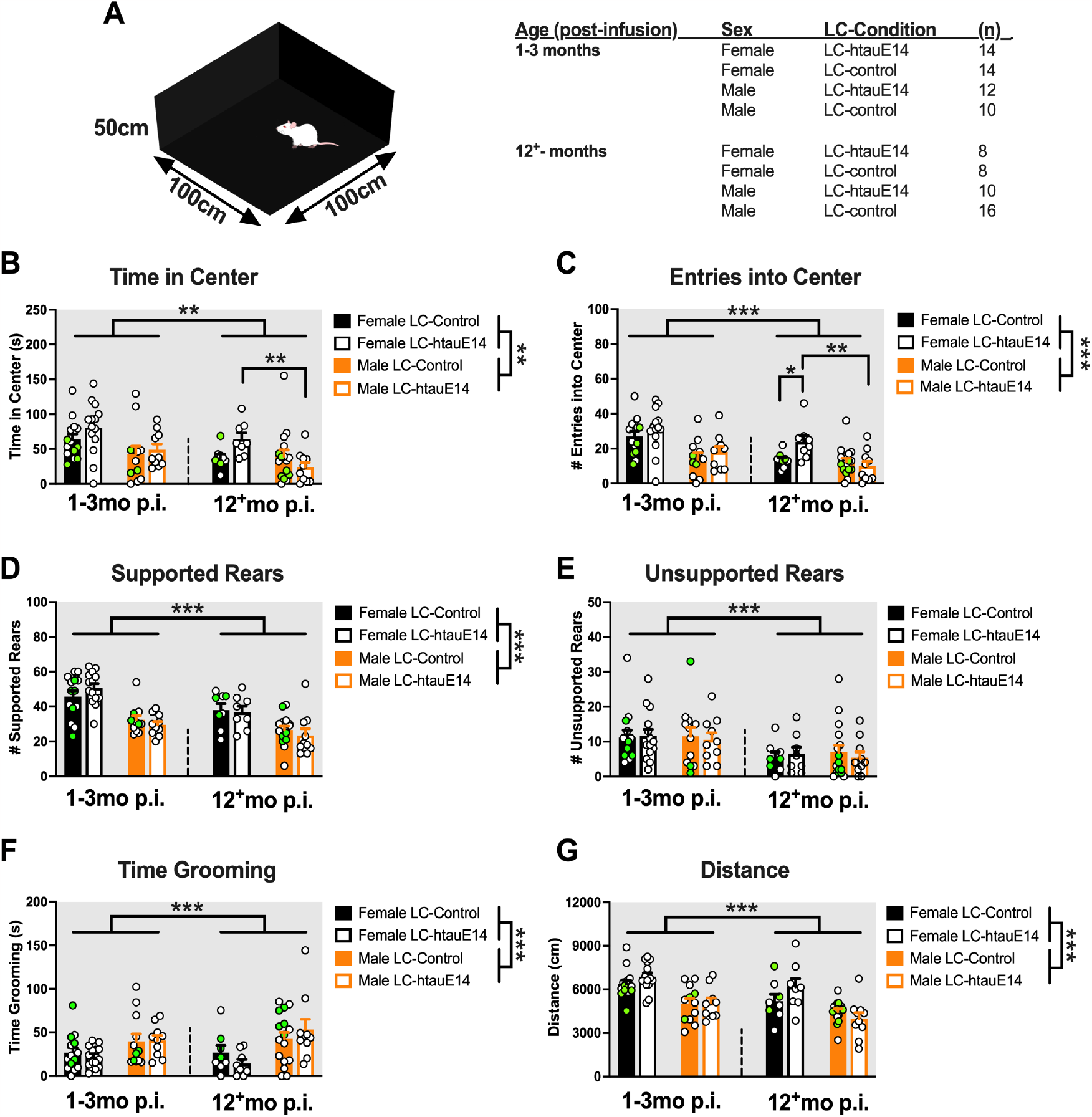
The effects of LC-htauE14 on open field (10min) behaviour in young and aged, male and female rats. **A**. Apparatus and list of subjects by condition. **B-C** Effects of LC-htauE14 infusion were observed in aged infused female rats in the center maze variables, increasing the time spent in center compared to aged LC-htauE14 infused male rats, and increased number of center entries compared to LC-Control female rats, and LC-htauE14 infused aged male rats. Main effects of age were observed across all measured variables (center, rearing, grooming and distance, **A-G**). Main effects of sex were also found in all variables with the exception of unsupported rears (**E**), with females demonstrating increased measures over males, except for time spent grooming (**F**). Green circles represent LC-AAV-GFP control infused rats. Data represent means ± s.e.m. **=min. p<0.01, ***min. p<0.001. Image MAW and SGW.

Effects of LC-htauE14 on anxiety measures in the open field maze: LC-htauE14 infusion most prominently affected measures thought to be reflective of anxiety, including the duration of time spent in the center of the maze, and the number of entries into the center in rats aged 12^+^mo post-infusion. The direction of these effects was dependent on sex. In the aged group there was a significant interaction of sex and LC-condition for the time spent in center variable (F_1,38_=5.316, p=0.0266). When explored, female LC-htauE14 rats spent more time (64.67±24.09s) in the center of the maze compared to male rats in the LC-htauE14 condition (23.87±22.52s; p=0.005). For the number of times the rats entered into the center of the maze, there was significant sex x LC-condition interaction in the aged animals (F_1,38_=5.8055, p=0.0209). Post hoc analyses revealed that aged female LC-htauE14 infused rats had a higher number of entries into the center (23.88±10.40) compared to female LC-Control rats of the same age (13.5±4.1; p=0.017) and aged female LC-htauE14 rats also had higher numbers of center entries compared to aged LC-htauE14 infused male rats (9.9±8.8; p<0.001).

#### Main effects of age on activity and anxiety in the open field maze

Younger (1-3mo p.i), rats had significantly higher scores on all variables compared to the 12^+^mo p.i. rats: Time in center (F_1,84_=6.64; p=0.01), number of center entries (F_1,84_=12.86; p=0.0006), supported (F_1,84_=11.09; p=0.001) and unsupported (F_1,84_=1.28; p=0.001) rears, time spent grooming (F_1,84_=18.83; p=0.00004) and distance traveled (F_1,84_=12.93; p=0.0005).

#### Main effects of sex on activity and anxiety in the open field maze

Similarly, main effects of sex were found across all variables save the number of unsupported rears. Male rats were found to spend less time in the center (F_1,84_=11.09; p=0.001), had fewer numbers of center entries (F_1,84_=11.09; p=0.00001), fewer unsupported rears (F_1,84_=49.61; p=0.000000) and traveled less distance than female rats (F_1,84_=38.73; p=0.000000) during the session. Males did spend more time grooming than females (F_1,84_=14.67; p=0.0002) and a sex x age interaction was also found for the grooming variable (F_1,84_=5.56; p=0.023).

### Simple and Difficult Odour Discrimination

To determine age and sex-dependent changes in odour discrimination tasks in LC-htauE14 infused rats, we employed two odour discrimination tasks using two sets of odourants. In the simple discrimination task (SOD) rats were exposed to two distinct odourants (banana and almond extract), and in a second difficult test (DOD), chemically similar odourants (heptanol 0.001% versus 50:50 0.001% heptanol/octanol) were examined.

#### Simple Odour Discrimination (SOD)

Two variables were examined in the odour discrimination tests: percentage of times the rat nose poked the target sponge as first choice, and the latency to nose poke the target odour sponge (Figure 2). In the SOD, no differences were found in either young or old LC-htauE14 infused rats to nose poke the correct sponge. An examination of the mean latency to nose poke across trials a significant age, sex and LC-condition interaction (F_1,63_=4.6569; p=0.03); further posthoc analysis determined the aged LC-htauE14 infused male rats took longer to nosepoke the correct target sponge (99.38±21.54s) than male rats in the aged LC-Control group (68.0±28.64s; p=0.03). The aged male LC-htauE14 infused rats also took longer to choose the correct target sponge than aged LC-htauE14 infused female rats (72.18±23.2s; p=0.04), and the younger male LC-htauE14 rats tested 1-3mo post-LC infusion (63.03±28.67; p=0.001).

**Figure 2.**
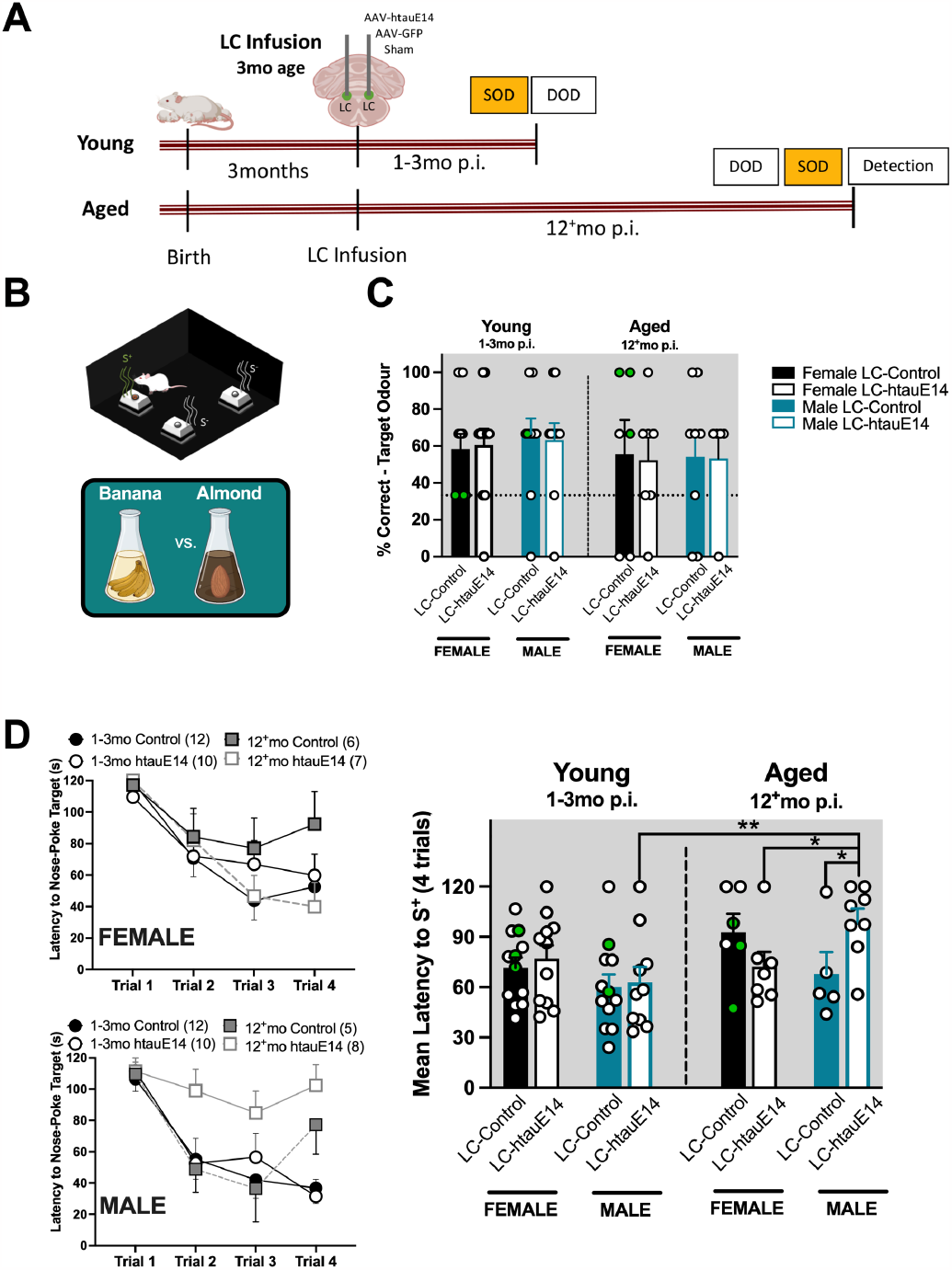
The effects of LC-htauE14 on a simple (distinct) odour discrimination (SOD) task in young and aged, male and female rats. **A**. Experimental timeline for young and aged rats highlighting SOD test (orange). **B**. Apparati (open field) and three sponges baited with S^-^(2) or S^+^(1) odours. Simple (distinct) odours were banana and almond almond extract (counterbalanced). **C**. Percent nosepoking target S^+^ sponge as first choice demonstrated there were no differences across age, sex or LC condition. **D**. Latency to nosepoke the target sponge across trials (x-y plot). The mean latency to nosepoke the target sponge was greatest in aged, male LC-htauE14 infused rats when compared to LC-Control male rats of same age, female LC-htauE14 infused female rats of the same age, and young male LC-htauE14 infused rats. Green circles represent LC-AAV-GFP control infused rats. Data represent means ± s.e.m. *=min. p<0.05, **min. p<0.01. Images MAW, SGW, Biorender.

#### Difficult Odour Discrimination (DOD)

When the more chemically similar odourants (heptanol 0.001% versus 50:50 0.001% heptanol/octanol) were examined in the DOD, a significant age and LC-group interaction was found in the percentage of time rats chose the correct odour infused sponge (F_1,63_=11.37; p=0.001; Figure 3). Further analysis found differences in LC-htauE14 and LC-Control rats within each sex (p=0.047 females, and p=0.027 in the male groups). Analysis of the latency to choose the correct odoured sponge found a significant interaction of age, sex and LC-group (F_1,64_=9.3888; p=0.003). Ultimately, it was shown that the aged male LC-htauE14 infused rats took significantly longer to choose the target sponge (95.94±27.01s) than aged male LC-Control infused rats (35.75±26.39s; p<0.00006), aged female LC-htauE14 infused rats (p<0.00006) and young LC-htauE14 infused male (p<0.00005) and female rats (p<0.000006).

**Figure 3.**
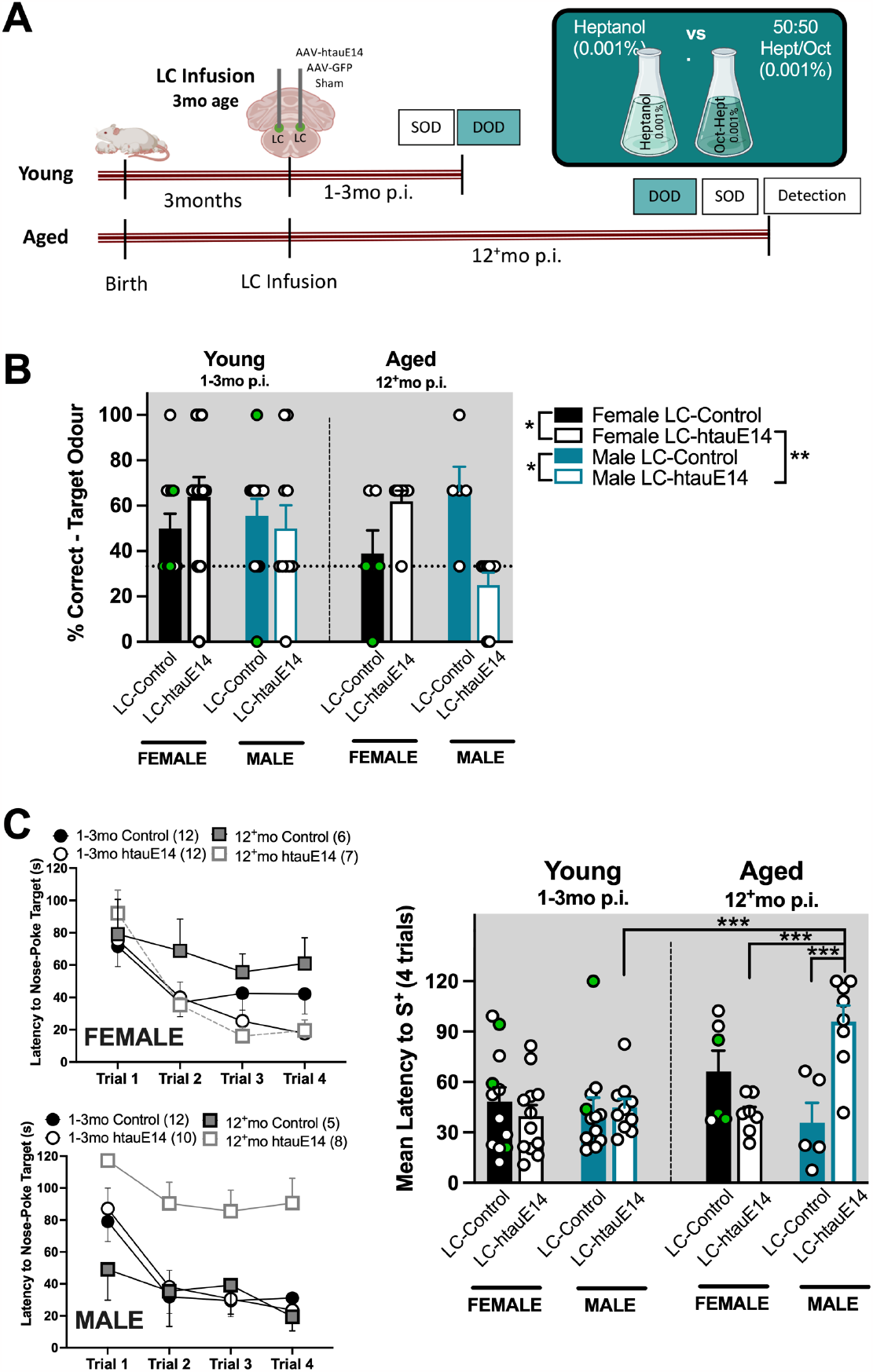
The effects of LC-htauE14 on a similar odour discrimination (DOD) task in young and aged, male and female rats. **A**. Experimental timeline for young and aged rats highlighting SOD test (teal), and difficult odour pair (0.001% heptanol and 50:50 of 0.001% heptanol and 0.001% octanol. **B**. Analysis of the percent nosepoking target S^+^ sponge as first choice found sex divergent effects in the LC-htauE14 conditions; LC-htauE14 male rats, had lower percent correct choices than male LC-Control infused rats, and female LC-htauE14 infused rats. Female LC-htauE14 infused rats unpredictably had higher percent correct choices than female rats in the LC-Control conditions. **C**. Latency to nosepoke the target sponge across trials (x-y plot) in DOD test. The mean latency to nosepoke the target sponge. Aged male LC-htauE14 infused rats took longer to nosepoke the target sponge. These results, taken together suggest impairment specific to aged LC-htauE14 infused male rats in the DOD test. Green circles represent LC-AAV-GFP control infused rats. Data represent means ± s.e.m. *=min. p<0.05, **min. p<0.01, ***min. p<0.001. Images Biorender.

#### Odour Detection: Simple and Difficult Odour Detection in Aged Male and Female LC-htauE14 Rats

Due to the diminished performance of aged male LC-htauE14 infused rats in the simple and difficult odour discrimination tests, both aged female and male rats were exposed to a set of habituation-dishabituation trials to examine if the impairment involved a decreased ability to detect the odours (Figure 4). Three 5min habituation trials (Trials 1-3) to two identical sponges (A and B) both baited with mineral oil only were presented, followed by a fourth Novel trial in which Sponge A was replaced with an identical sponge with corners baited with either of the simple odours (Simple Odour Detection), or either of the two difficult odours (Difficult Odour Detection). Student t-tests were performed on the percentage of time spent with Sponge A during the novel odour trial (4^th^ or dishabituation trial) and the third trial of the mineral oil habituation test. For the aged LC-htauE14 male rats, responses to the novel odours were not significantly different from the respective Trial 3 in both the simple and difficult detection tests (p>0.05), suggesting that the rats were not able to detect the change when novel odourants were presented after the mineral oil trials. This deficit was more pronounced in the difficult odour conditions suggesting that aged LC-htauE14 infused male rats were more selectively impaired in their ability to detect the difficult novel odours.

**Figure 4.**
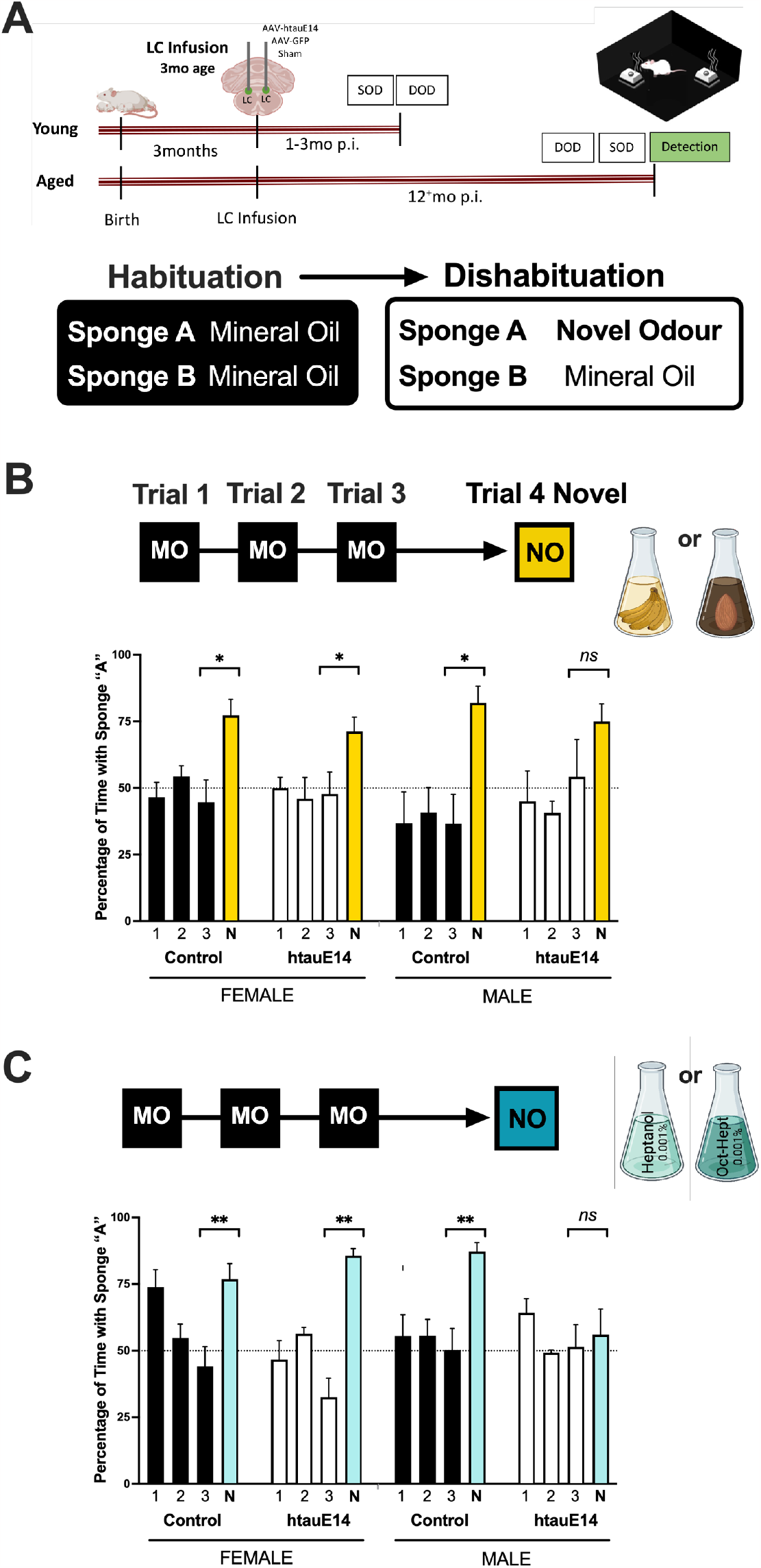
The effects of LC-htauE14 on a simple (distinct) and difficult (similar) odour detection in aged, male and female rats. **A**. As aged male LC-htauE14 infused rats appeared impaired in both the SOD and DOD tests, aged rats (male and female) were exposed to habituation-dishabituation odour detection tests. In Trials 1-3 rats were exposed to two sponges impregnated with mineral oil (MO) only (habituation). On the fourth trial one of the two sponges contained the novel odour (dishabituation). This test was performed once for each of the odour pairs (SOD, and DOD). **B**. Results of habituation-dishabituation test for SOD odours (banana or almond). Across all LC-conditions, ages and sexes, all rats spent a greater percentage time with the novel odour sponge in Trial 4 compared to the last habituation trial (Trial 3), although these results were not significant in the aged, male LC-htauE14 group **C**. Results of habituation-dishabituation test for DOD odours (heptanol or heptanol/octanol). Across all LC-conditions with the exception of the aged, male LC-htauE14 infused group, spent a greater percentage time with the novel odour sponge in trial 4 compared to the last habituation trial (Trial 3). These results suggest that aged male LC-htauE14 infused rats are moderately impaired at detecting the SOD odourants and more severely impaired at detecting the DOD odourants. Data represent means ± s.e.m. *=min. p<0.05, **min. p<0.01. n=5 all groups.

### Effects of htauE14 on Locus Coeruleus Single Unit Response in Aged Rat

Neuron counts and area measures of LC Nissl-stained sections did not reveal differences in the average numbers of neurons or differences per section in LC-htauE14 infused rats compared to LC-Control rats (214.3±4.045 and 215.8±19.77, respectively), or LC area measures (0.053±0.0085mm^2^ and 0.057±0.0041mm^2^, respectively).

Putative single LC units were recorded in behaviourally naïve, aged, male LC-htauE14 (n=4), and LC-Control (n=3) rats. Parameters of the single units were analyzed and there were no differences in waveform amplitude (LC-htauE14, 54.2μV±14.4; LC-Control, 50.6μV±7.55; t(5)=0.3915; p=0.712), or slope (LC-htauE14, -113.16μV/ms±47.3; LC-Control, -113.4μV/ms ±25.18; t(5)=-0.0073; p=0.994); however, measures of waveform half-width trended more closely to significance (LC-htauE14, 492.0μs±24.9; LC-Control, 551.3μs±37.96; t(5)=-2.3820; p=0.063).

Measures of baseline firing frequency were sampled for 60 s and organized in 1-s bins. Analysis of firing frequency from the 60-s epoch found the basal firing rate of recorded LC neurons was higher in LC-htauE14 infused rats (2.30Hz±0.534) compared to age-matched LC-Control rats (0.945Hz±0.234; t(5)=4.04; p=0.01; see Figure 5F). A non-linear sine wave function was applied revealing an increase in sine wave amplitude (increased neuronal firing frequency) and decreased sine wave length (increased oscillation frequency) in LC-htauE14 infused rats compared to rats in the LC-Control condition (Figure 5G).

**Figure 5.**
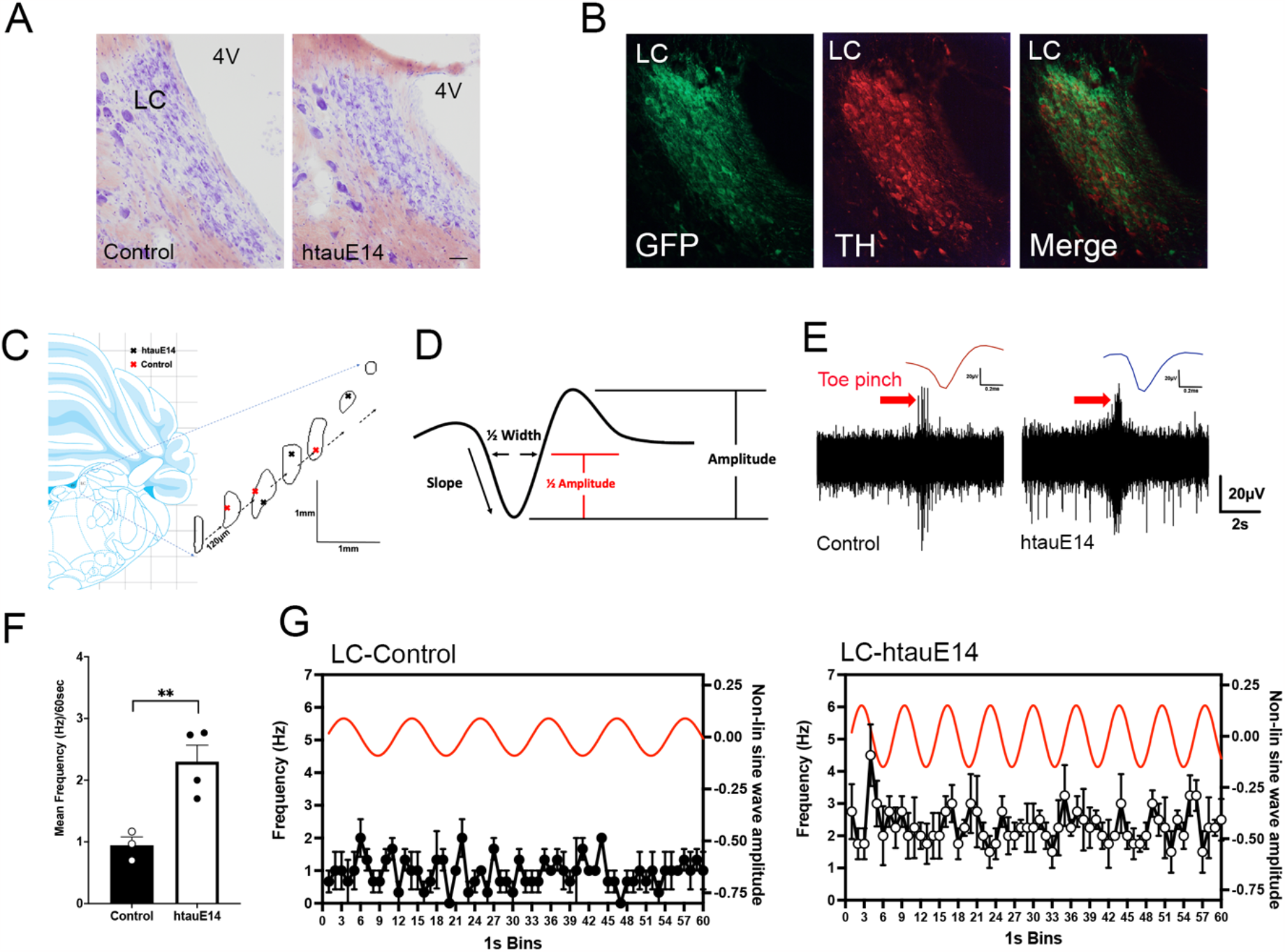
The effects of LC-htauE14 on LC single unit recording in aged male rats. **A**. Sample Nissl-stained sections in Control and LC-htauE14 infused rats. No differences in area measures or LC neuron counts were found across conditions. Scale is 50µm in both A and B. **B**. Co-labelling of htauE14-GFP (green), and tyrosine hydroxylase (red) show selective co-localization in LC cells. **C**. Recording electrode placements in Control (red cross) and LC-htauE14 (black cross) infused rats (image adapted from Paxinos and Watson, 1998). **D**. Sample LC unit with parameters used to identify and sort units. **E**. Sample LC units and response to contralateral toe-pinch stimuli in Control and LC-htauE14 infused rats. **F**. Mean frequency of LC baseline activity (60s) revealed an increased baseline frequency firing rate in LC-htauE14 infused rats compared control rats. **G**. A non-linear sine wave function was applied to the entire baseline frequency variable over the 60s recording revealing an increased oscillatory frequency and higher overall amplitude in LC-htauE14 infused rats compared to the sine wave function of the control rats. Data represent means ± s.e.m.; **min. p<0.01.

### Analysis of pre- and post-synaptic markers synaptophysin and PSD-95 labelling in piriform cortex (PCx) after LC-htauE14 expression

Brightfield, DAB-metal PSD95 and synaptophysin immunostained sections were analyzed for relative density staining in PCx, and converted to the average ROD levels of LC-Control levels in each condition (see Figure 6). Examination of the postsynaptic marker PSD95 did not reveal differences in the ROD due to LC-htauE14 infusion; however, main effects of age (F_1,27_=8.80; p=0.006) and sex (F_1,27_=4.34; p=0.04) were found (Figure 6). A sex and LC-condition interaction was also found, however posthoc analysis revealed that these measures were higher in LC-Control female rats compared to LC-Control male rats (p=0.009).

**Figure 6.**
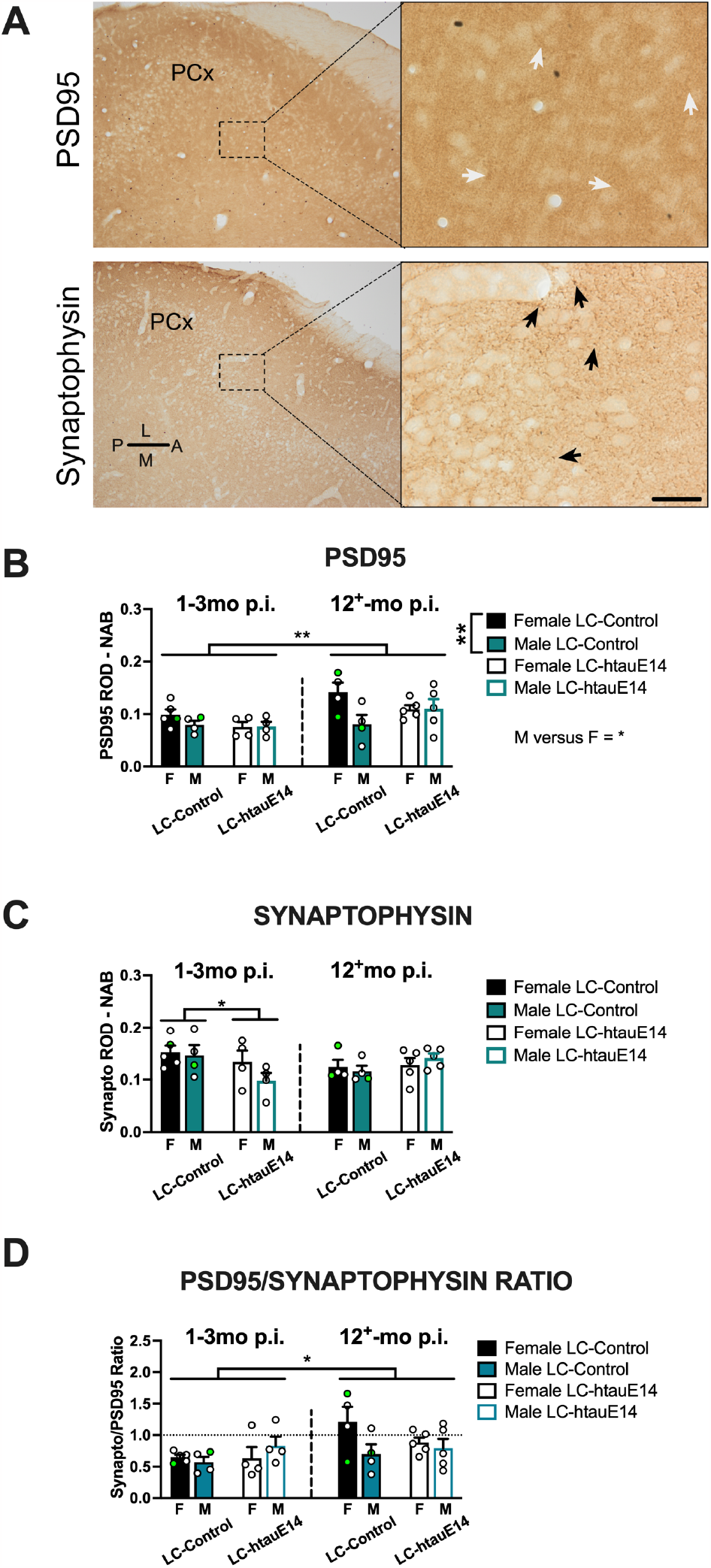
Analysis of postsynaptic (PSD95) and presynaptic (synaptophysin) markers in piriform cortex (PCx) of young and aged, male and female LC-htauE14 infused rats. **A**. Samples of PSD95 and synaptophysin immunostaining and ROD region sampled in tissue sectioned in the horizontal plane, and magnified for detail (right panels). Arrows identify positively stained PSD95 puncta (white) and synaptophysin (black). Anatomical direction represented in PSD95 image. Scale is 50µm in right panels and 300µm in left panels. **B**. Mean raw ROD measures less ROD of no-1 antibody (NAB) in PSD95 immunostained sections. No significant effects of LC-htauE14 was found, however there was a significant main effect of age (aged > young), and sex (female > male) and a significant age x LC-condition found (female LC-Control > male LC-Control). **C**. Mean raw ROD measures less ROD NAB in synaptophysin labelled sections. LC-htauE14 decreased synaptophysin ROD in rats when measured at the early time period (1-3mo p.i.). This difference was transient as it was not evident in the aged rats. **D**. Analysis of the PSD95/synaptophysin ROD ratio measure found only an effect of age with aged rats expressing higher ratio measures compared to ratio measures in young rats. *min. p<0.05, **min. p<0.01.

For the presynaptic marker synaptophysin, an age by LC-condition interaction was also found (F_1,27_=5.29; p=0.03), and further analysis identified a significantly lower staining density in male and female LC-htauE14 rats 1-3mo post-infusion compared to rats in the LC-Control condition at this same age. This decrease relative to LC-Control staining was not observed in the aged rat groups (Figure 6C). When raw ROD measures of PSD95 and synaptophysin were converted to a ratio measure, differences were found as a function of age (F_1,27_=5.20; p=0.03) with the ratio being higher in aged rats compared to younger adult rats, an effect that was not dependent upon LC-condition, or sex (Figure 6D)

## DISCUSSION

We examined behavioural measures of activity and anxiety in an open field maze, and tests of simple and difficult odour discrimination in young (1-3mo p.i.) and aged (12^+^mo p.i.) male and female rats in a LC-htauE14 rat model of pretangle stage tau dysfunction. In an open field test, aged rats had lower overall measures for all variables than the younger counterparts, while aged female LC-htauE14 infused rats demonstrated lower measures of behavioural inhibition to novel environments (time in center variable) compared to LC-Control infused and aged male htauE14-infused rats. In the odour tests, aged male LC-htauE14 rats were observed to be impaired in some measures within the simple (dissimilar) odour pairs; however, during the difficult odour discrimination test with chemically similar stimuli, aged male LC-htauE14 infused rats could not detect the presentation of a novel difficult odourant in habituation/dishabituation odour test. Single cell recordings of LC neurons in aged, behaviourally naïve, male rats revealed a significantly higher level of baseline firing rate of identified LC neurons htauE14 infused rats compared to LC neurons in rats in the LC-Control group. Additionally, it was found when examined on a macroscale (60s) that aged male LC-htau14 rats had increased sine wave power and increased variability of frequency (amplitude) when contrasted with that of the control rats. Finally, densitometric analyses of pre- and post-synaptic markers, synaptophysin and PSD95 in the primary olfactory cortex (PCx) found only transient decreases in synaptophysin percentage density measures in LC-htauE14 infused rats 1-3mo p.i., which were within levels measured in control rats when examined 12^+^mo post infusion. Synaptophysin/PSD-95 density ratios were higher in aged rats (both sexes) compared to rats examined at the younger age, regardless of LC condition. Results are discussed in reference to the effects of LC-htauE14 on behavioural and physiological changes, and in reference to age and sex, where appropriate.

### The effects of age, sex and LC-htauE14 on measures of activity and anxiety in the OFT

As has been reported previously, we found overall main effects of age and sex on measures of activity and anxiety in the open field. Previous reports across rat strains typically identify increased levels of activity in female rats compared to age-matched male rats (e.g., Mancini et al., 1989; Belviranli et al., 2012). Lower levels of activity have also been reported in aged male rats (Schulz et al., 2007; Hamezah et al., 2017), with fewer studies examining activity and anxiety in the open field as a function of both age and sex (see Belviranli et al., 2012 for exception).

Infusion of htauE14 into the LC most significantly affected the center of maze variables (time in center, and entries into center) selective to the aged (12^+^mo p.i.) category. These effects were dependent on sex, with female LC-htauE14 infused rats spending more time in the center of the maze compared to male LC-htauE14 infused rats. Similarly, female LC-htauE14 infused rats entered the center more frequently than male LC-htauE14 infused rats, and female rats in the LC-Control condition. These results were not activity-dependent changes as measured by distance or other activity related variables, and would suggest that aged female LC-htauE14 had lower levels of anxiety. In a similar series of experiments examining activity and anxiety in the open field test, Omoluabi et al. (2021) reported that LC-htauE14 (at 9-10mo p.i.) did not significantly alter anxious behaviour in the open field test i.e., percentage of time spent freezing, however in this circumstance maze center-focused variables, or sex-related comparisons were not included in the analysis. Similar to the present study, they did not find that LC-htauE14 infusion affected the percentage of time spent rearing, or the distance travelled during the 10-min test.

Lower anxiety scores in the open field test have been reported in AD animals using other rodent models of AD e.g., 5xFAD mice (Forner et al., 2021; Miao et al., 2023), so it may be that the aged female LC-htauE14 infused rats fit a previously reported phenotypic AD profile (for counterexamples in AD rat models see Galeano et al., 2014 for results in aged male rats and Srivastava et al., 2023 for transient sex and age-dependent measures in the open field). It should be noted that LC-htauE14 may also affect open field behaviour through other cognitive and behavioural systems. As examples, LC-norepinephrine projections in forebrain regions are known to influence novelty exploration (e.g. Sara et al., 1995), fear/anxiety (e.g. Gu et al., 2020), spatial/contextual memory (e.g. Grella et al., 2019), decision making (e.g. Rodberg et al., 2023), and attentional processes (e.g. Aston-Jones et al., 1991) all of which are systems known to be modified in individuals living with AD and related dementias. It is not possible to recognize with certainty which of these systems LC-htauE14 may be influencing in the open field test during this pretangle stage. It would be of benefit to include further behavioural, age- and sex-dependent changes using physiological and anatomical assays targeting other LC-modulated behavioural responses in LC-htauE14 model.

### The age- and sex dependent effects of early adult LC-htauE14 on olfactory discrimination and detection in aging male and female rats

In this study, we report that LC-htauE14 infusion in 3mo old rats significantly impaired olfactory detection and discrimination of similar odour pairs selectively in aged male rats, while discrimination and detection of the odourants remained intact in female LC-htauE14 infused rats. These results support, in part, the results of previous studies examining LC-htauE14 in TH-Cre^+/-^ rats infused at similar time points. Both Ghosh et al. (2019) and Omoluabi et al. (2021) report impairment of the dilute similar odour pairs (0.001% heptanol and a 50:50 solution of heptanol/octanol of same concentration) during discrimination/detection in LC-htauE14 male and female rats when data were combined for both sexes. In the present study, both sex- and age-dependent effects of LC-htauE14 on odour detection and discrimination of the similar odourants were observed and were most prominent in aged male LC-htauE14 infused rats. A future consideration would be to examine discrimination and detection of the simple odours (banana and almond) at the same dilution as was examined in the heptanol/octanol tests to determine if this is a generalized olfactory deficit in aged LC-htauE14 male rats. These deficits do not appear to be the result of changes in pre-or post-synaptic density changes in the PCx, as indexed by synaptophysin and PSD-95, respectively. The earlier studies detail loss of immunostained fibres positive for the norepinephrine synthesizing enzyme dopamine-β-hydroxylase (DβH), and norepinephrine transporters (NET) in PCx (Ghosh et al., 2019 and Omoluabi et al., 2021) and the dentate gyrus (Omoluabi et al., 2021), two structures known to be essential for olfactory pattern separation and discrimination learning (Shakhawat et al., 2015; Woods et al., 2020). Alterations in LC-norepinephrine input (Bouret and Sara, 2002) and the LC-ventral tegmental area (VTA) dopaminergic influence (Ghosh et al., 2021) in PCx have been associated with odour discrimination learning and may contribute to the impairment observed in aged male LC-htauE14 rats. The failure of the aged male rats to detect the dilute chemicaly similar odours in the habituation-dishabituation test also suggests additional sex-dependent neuropathological changes within this system that may be related to LC-norepinephrine (e.g., DβH/NET), not yet examined in relation to sex, or may be the consequence of other LC-htauE14-related neuropathological changes related to the organizational or activational influences of steroid sex hormones.

The LC-htauE14 model is characterized by LC-originated spread of htauE14-GFP to entorhinal cortex (in preparation). In rodents, the lateral and medial entorhinal cortices receive olfactory input from the olfactory bulb (Kosel et al., 1981), anterior olfactory nucleus (Haberly and Price, 1978), and the PCx with the estimate that the PCx contributes one-third of the total afferent input into both entorhinal regions (Burwell and Amaral, 1998). The medial entorhinal cortex is also a prominent contributor to the parahippocampal and hippocampal system mapping spatial representations (see Moser et al., 2017 for review). Amani et al. (2021) report impaired plasticity (tetanic long-term potentiation; LTP) at both the lateral and medial perforant path-dentate gyrus synapse in hippocampal slices of male Long-Evans rats, at 8-10mo of age, an effect that was not dependent on loss of the presynaptic vesicular protein, synaptophysin at the synapse in the outer molecular layer of dentate gyrus. Recent in vivo work by Walling et al. (2023) also described an impairment of medial perforant path (MPP)-dentate gyrus frequency-induced LTP in aged (18-20mo old) male Sprague-Dawley rats, where plasticity of the MPP evoked population spike in dentate gyrus was significantly lower in aged male rats compared to younger adult male rats (5-9mo old), and also impaired when contrasted with aged female rats. Although, the age-dependent impairment of LPP-dentate gyrus LTP in male rats has yet to be examined in female rats, weakened plasticity of olfactory input from the LPP/MPP selective to aged male LC-htauE14 infused rats may make aged male rats more susceptible to learning and memory deficits in tasks that are dentate gyrus-dependent, such as odour discrimination.

### The effects of early adult LC-htauE14 on the physiological responses of LC neurons

As aged male rats were impaired in odour discrimination and detection tests, we recorded putative single unit LC neurons in behaviourally naïve LC-htauE14, and LC-Control aged male rats. Single unit LC neurons from LC-htauE14 infused rats were found to have elevated rates of baseline firing (hyperactivity) compared to rats in the LC-Control condition. Sine wave analysis of firing rate (60s) also revealed an overall increase in the non-linear sine wave amplitude suggesting larger overall variations in LC-htauE14 firing rate. Shorter sine wavelength cycles were also found in LC neurons of htauE14 infused rats, which would suggest faster phases of basal oscillatory activity. In the present study, the baseline firing rate of the putative LC neurons of both LC-htauE14 and LC-Control groups were within the range previously reported in urethane anesthetized aged male rats (e.g. Shirokawa et al., 2000), although the firing rates in the LC-htauE14 infused rats were consistently higher than LC-Control baseline rates. Previous in vitro and in vivo studies examining changes in LC firing and action potential parameters after selective noradrenergic or catecholaminergic chemical lesions indicate increased basal firing rates, changes in action potential duration, and alterations in Ca^2+^ currents; all of which were postulated to be compensatory responses to the catecholaminergic lesions (Chiodo et al., 1983; Magnuson et al., 1993). Similarly, we report near significant reductions in LC action potential half-width measures in the small number of LC-htauE14 infused rats, and subtle (when compared to DSP-4 lesioned animals), but consistent, increases in baseline firing rates. These results, in addition to previously reported decreases in LC noradrenergic input with concomitant increase in β_1−_adrenergic receptor density in the PCx of LC-htauE14 infused rats (Ghosh et al., 2019) suggest the responses in the htauE14^+^ LC neurons may be compensatory to pretangle stage tau accumulation in these cells.

The only other in vivo study examining physiological responses of LC neurons with confirmed pretangle tau pathology was performed recently in Tg F344-AD rats (Kelberman et al., 2023) in which accumulation of persistently phosphorylated tau is confirmed at 6mo of age (see Rorabaugh et al., 2017). Kelberman et al. reported hypoactivity of LC neuron baseline firing in chloral hydrate anesthetized male and female rats at 6 and 15mo age. In the Tg F344-AD rats, a distinction was observed in the phasic responses of these neurons to footshock where 6mo old rats exhibited an elevated phasic response to footshock, while the LC neurons of the aged rats had a muted response (hypoactivity). Functional responses to sensorineural stimulation were not directly quantified in the present study, although LC phasic responses to contralateral toe pinch were elicited in both LC-htauE14^+^ and LC-Control rats. It is possible that evoked phasic LC activity in aged LC-htauE14 rats may be elevated over the response of LC neurons in control rats, similar to the increased tonic basal activity reported here. Equally plausible however, LC responses to evoked phasic activation in LC-htauE14 rats may be hypoactive, similar to the results in the Kelberman Tg F344-AD study, despite being hyperactive under tonic baseline firing conditions. It would be of great interest to examine the physiological responses of LC neurons in both models under the influence of functional and evoked, phasic and tonic stimulation as chemogenetic manipulation of LC neurons in Tg F344-AD rats (Rorabaugh et al., 2017), and phasic, but not tonic optogenetic stimulation in the LC-htauE14 model (Omoluabi et al., 2021) rescued behavioural impairments in these models.

In conclusion, we found age- and sex-dependent changes in behaviour and LC physiology in a rat model modelling Braak’s pretangle stages of LC-originated tau dysfunction. The results of this study found that the influence of LC-htauE14 is expressed differentially in male and female rats, further identifying the importance and relevance of including sex as a specific variable of interest in rodent models of AD. To fully characterize pretangle stage dysfunction, the age- and sex-dependent effects of LC-htauE14 on LC activity (basal, phasic and tonic) related to changes in cognition and behaviour warrants further exploration.

## Author Contributions

SGW and CWH designed research. ATB, EAC, ODED, ATJ, and MAW performed research; ATB, EAC, ODED, ATJ, RV, and MAW analyzed data (alphabetical order); SGW, DMS, ODED, ATB, and MAW wrote the paper. All authors reviewed the manuscript and approved the submitted version.

## Acknowledgements

The authors would like to thank the many individuals who contributed to portions of this project and would like to express gratitude to all individuals who participated in the daily care and feeding schedules of the animal colony. Thank you also to Dr. Qi Yuan for comments on this manuscript.

